# Single-cell transcriptomic analysis of SARS-CoV-2 reactive CD4^+^ T cells

**DOI:** 10.1101/2020.06.12.148916

**Authors:** Benjamin J. Meckiff, Ciro Ramírez-Suástegui, Vicente Fajardo, Serena J Chee, Anthony Kusnadi, Hayley Simon, Alba Grifoni, Emanuela Pelosi, Daniela Weiskopf, Alessandro Sette, Ferhat Ay, Grégory Seumois, Christian H Ottensmeier, Pandurangan Vijayanand

## Abstract

The contribution of CD4^+^ T cells to protective or pathogenic immune responses to SARS-CoV-2 infection remains unknown. Here, we present large-scale single-cell transcriptomic analysis of viral antigen-reactive CD4^+^ T cells from 32 COVID-19 patients. In patients with severe disease compared to mild disease, we found increased proportions of cytotoxic follicular helper (T_FH_) cells and cytotoxic T helper cells (CD4-CTLs) responding to SARS-CoV-2, and reduced proportion of SARS-CoV-2 reactive regulatory T cells. Importantly, the CD4-CTLs were highly enriched for the expression of transcripts encoding chemokines that are involved in the recruitment of myeloid cells and dendritic cells to the sites of viral infection. Polyfunctional T helper (T_H_)1 cells and T_H_17 cell subsets were underrepresented in the repertoire of SARS-CoV-2-reactive CD4^+^ T cells compared to influenza-reactive CD4^+^ T cells. Together, our analyses provide so far unprecedented insights into the gene expression patterns of SARS-CoV-2 reactive CD4^+^ T cells in distinct disease severities.

## INTRODUCTION

Coronavirus disease 2019 (COVID-19) is causing substantial mortality, morbidity and economic losses (Nicolas Vabret et al., 2020; Tay et al., 2020) and effective vaccines and therapeutics may take several months or years to become available. A substantial number of patients become life-threateningly ill, and the mechanisms responsible for causing severe respiratory distress syndrome (SARS) in COVID-19 are not well understood. Therefore, there is an urgent need to understand the key players driving protective and pathogenic immune responses in COVID-19 (Nicolas Vabret et al., 2020). This knowledge may help devise better therapeutics and vaccines for tackling the current pandemic. CD4^+^ T cells are key orchestrators of anti-viral immune responses, either by enhancing the effector functions of other immune cell types like cytotoxic CD8^+^ T cells, NK cells and B cells or through direct killing of infected cells (Sallusto, 2016). Recent studies in patients with COVID-19 have verified the presence of CD4^+^ T cells that are reactive to SARS-CoV-2 (Braun et al., 2020; Grifoni et al., 2020; Thieme et al., 2020). However, the nature and types of CD4^+^ T cell subsets that respond to SARS-CoV-2 and whether they play an important role in driving protective or pathogenic immune responses remain elusive. Here, we have analyzed single-cell transcriptomes of virus-reactive CD4^+^ T cells to determine associations with severity of COVID-19 illness, and to compare the molecular properties of SARS-CoV-2-reactive CD4^+^ T cells to other common respiratory virus-reactive CD4^+^ T cells from healthy control subjects.

## RESULTS

### CD4^+^ T cell responses in COVID-19 illness

To capture CD4^+^ T cells responding to SARS-CoV-2 in patients with COVID-19 illness, we employed the antigen-reactive T cell enrichment (ARTE) assay (Bacher et al., 2016; Bacher et al., 2019; Bacher et al., 2013) that relies on *in vitro* stimulation of peripheral blood mononuclear cells (PBMCs) for 6 hours with overlapping peptide pools targeting the immunogenic domains of the spike and membrane protein of SARS-CoV-2 (see **STAR Methods** (Thieme et al., 2020)). Following *in vitro* stimulation, SARS-CoV-2-reactive CD4^+^ memory T cells were isolated based on the expression of cell surface markers (CD154 and CD69) that reflect recent engagement of the T cell receptor (TCR) by cognate major histocompatibility complex (MHC)-peptide complexes (**Figure S1A**). In the context of acute COVID-19 illness, CD4^+^ T cells expressing activation markers have been reported in the blood (Braun et al., 2020; Thevarajan et al., 2020); such CD4^+^ T cells, presumably activated *in vivo* by endogenous SARS-CoV-2 viral antigens, were also captured during the ARTE assay, thereby enabling us to study a comprehensive array of CD4^+^ T cell subsets responding to SARS-CoV-2. We sorted > 200,000 SARS-CoV-2-reactive CD4^+^ T cells from > 1.3 billion PBMCs isolated from a total of 32 patients with COVID-19 illness (22 hospitalized patients with severe illness, 9 of whom required intensive care unit (ICU) treatment, and 10 non-hospitalized subjects with relatively milder disease, **Figures 1A, 1B** and **Table S1**). In addition to expressing CD154 and CD69, sorted SARS-CoV-2-reactive CD4^+^ T cells co-expressed other activation-related cell surface markers like CD38, CD137 (4-1BB), CD279 (PD-1) and HLA-DR (**Figures 1C, S1B** and **Table S2**).

**Figure 1.**
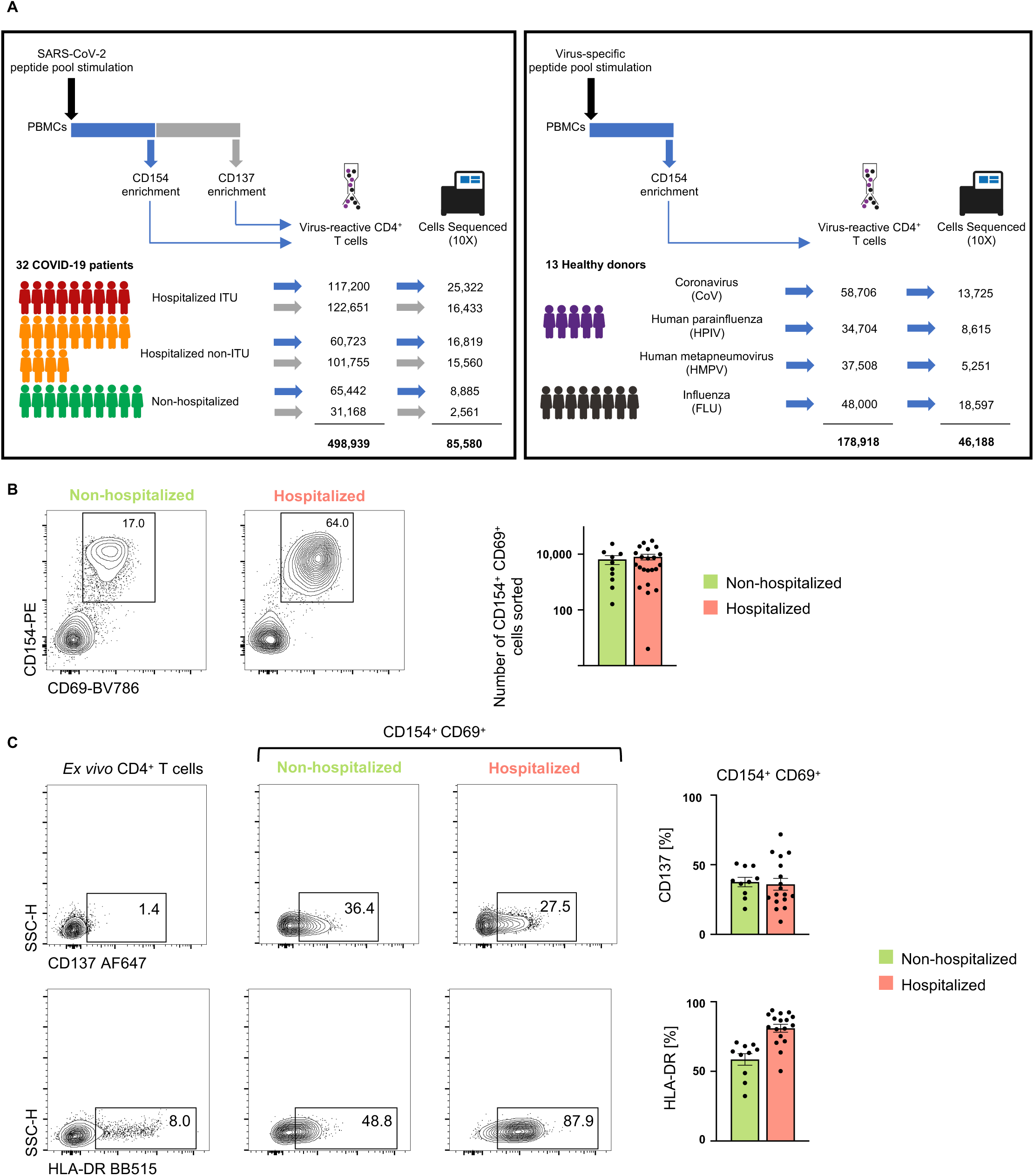
CD4^**+**^ T cell responses in COVID-19 illness. **(A)** Study overview. **(B)** Representative FACS plots showing surface staining of CD154 (CD40L) and CD69 in memory CD4^+^ T cells stimulated for 6 hours with SARS-CoV-2 peptide pools, post-enrichment (CD154-based), in hospitalized and non-hospitalized COVID-19 patients (left), and summary of number of cells sorted (right); Data are mean +/-S.E.M. **(C)** Representative FACS plots (left) showing surface expression of CD137 (4-1BB) and HLA-DR in memory CD4^+^ T cells *ex vivo* (without *in vitro* stimulation) and in CD154^+^ CD69^+^ memory CD4^+^ T cells following stimulation, post-enrichment (CD154-based). (Right) Percentage of CD154^+^ CD69^+^ memory CD4^+^ T cells expressing CD137 (4-1BB) or HLA-DR in 17 hospitalized and 10 non-hospitalized COVID-19 patients; Data are mean +/-S.E.M.

Recent evidence from studies in non-exposed individuals (blood sample obtained pre-COVID-19 pandemic) indicates pre-existing SARS CoV2-reactive CD4^+^ T cells, possibly indicative of human coronavirus (HCoV) cross-reactivity. Such cells are observed in up to 50% of the subjects studied (Braun et al., 2020; Grifoni et al., 2020). To capture such SARS-CoV-2-reactive CD4^+^ T cells, likely to be coronavirus (CoV)-reactive, we screened healthy non-exposed subjects and isolated CD4^+^ T cells responding to SARS-CoV-2 peptide pools from 4 subjects with highest responder frequency (**Figures 1A** and **S1C**). Next, for defining the CD4^+^ T cell subsets and their properties that distinguish SARS-CoV-2-reactive cells from other common respiratory virus-reactive CD4^+^ T cells, we isolated CD4^+^ T cells responding to peptide pools specific to influenza (FLU) hemagglutinin protein (FLU-reactive cells, see **STAR Methods**) from 8 additional healthy subjects who provided blood samples before and/or after influenza vaccination (**Figure 1A** and **S1D**). CD4^+^ T cells responding to peptide pools specific to other common respiratory viruses like human parainfluenza (HPIV) and human metapneumovirus (HMPV) were also isolated from healthy subjects (**Figure S1C**). In total, we interrogated the transcriptome and T cells receptor (TCR) sequence of >100,000 viral-reactive CD4^+^ T cells from 45 subjects (**Figures 1A, S2A, S2B** and **Table S3**).

### SARS-CoV-2-reactive CD4^+^ T cells are enriched for T_FH_ cells and CD4-CTLs

Analysis of the single-cell transcriptomes of all viral-reactive CD4^+^ T cells from all subjects revealed 13 CD4^+^ T cell subsets that clustered distinctly, reflecting their unique transcriptional profiles (**Figures 2A-2D** and **Table S4**). Strikingly, a number of clusters were dominated by cells reactive to specific viruses (**Figures 2B** and **S2C**). For example, the vast majority of cells in clusters 1 and 10 were FLU-reactive (>65%), whereas cells in clusters 0,4,6,7 and 12 mainly consisted of SARS-CoV-2 reactive CD4^+^ T cells (>70%) from COVID-19 patients (**Figures 2B** and **S2C**). Conversely, cells in clusters (3,5 and 11) were not preferentially enriched for any given virus (**Figures 2B** and **S2C**). These findings suggest that distinct viral infections generate CD4^+^ T cell subsets with distinct transcriptional programs, although the timing of survey (acute illness *versus* past infection) might also contribute to their molecular states. Our data highlight substantial heterogeneity in the nature of CD4^+^ T cells generated in response to different viral infections on the one hand and shared features on the other.

**Figure 2:**
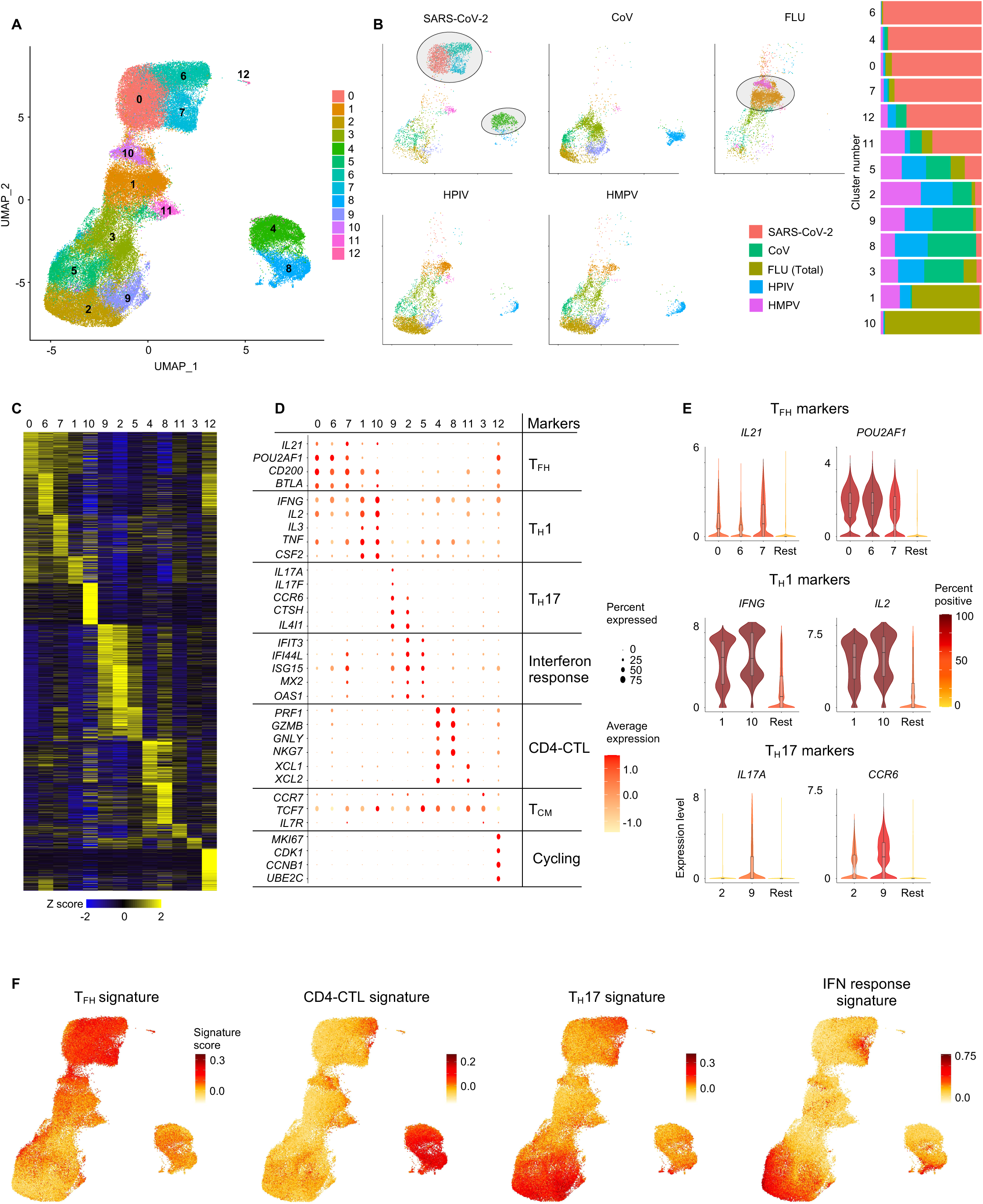
SARS-CoV-2-reactive CD4^+^ T cells are enriched for T_FH_ cells and CD4-CTLs. **(A)** Single-cell transcriptomes of sorted CD154^+^ CD69^+^ memory CD4^+^ T cells following 6 hours stimulation with virus-specific peptide megapools is displayed by manifold approximation and projection (UMAP). Seurat-based clustering of 91,140 cells colored based on cluster type. **(B)** UMAPs showing memory CD4^+^ T cells for individual virus-specific megapool stimulation conditions (left), and normalized proportion of each virus-reactive cells per cluster is shown (right). **(C)** Heatmap showing expression of the most significantly enriched transcripts in each cluster (see Table S4, Seurat marker gene analysis – comparison of cluster of interest *versus* all other cells, shown are the top 200 transcripts with adjusted P value < 0.05, fold change > 0.25 and >10% difference in the percentage of cells expressing differentially expressed transcript between two groups compared. **(D)** Plot shows average expression and percent expression of selected marker transcripts in each cluster. **(E)** Violin plots showing expression of T_FH_ (top), T_H_1 (middle) and T_H_17 (bottom) marker transcripts in designated clusters compared to an aggregation of remaining cells (Rest). Color indicates percentage of cells expressing indicated transcript. **(F)** UMAP showing T_FH_, CD4-CTL, T_H_17 and interferon (IFN) response signature scores for each cell.

The clusters enriched for FLU-reactive CD4^+^ T cells (clusters 1 and 10) displayed features suggestive of polyfunctional T_H_1 cells which have been associated with protective anti-viral immune responses (Seder et al., 2008). Such features include the expression of transcripts encoding for the canonical T_H_1 transcription factor T-bet, cytokines linked to polyfunctionality, IFN-*γ*, IL-2 and TNF, and several other cytokines and chemokines like IL-3, CSF2, IL-23A and CCL20 (**Figures 2D, 2E, S2E** and **S2F**). SARS-CoV-2-reactive CD4^+^ T cells were under-represented in these clusters (cluster 1 and 10, <2%), when compared to FLU-reactive cells (>70%) or HMPV-and HPIV-reactive cells (∼5-20%) (**Figure S2C**). Furthermore, SARS-CoV-2-reactive CD4^+^ T cells in cluster 1 expressed significantly lower levels of *IFNG* and *IL2* transcripts when compared to FLU-reactive cells (**Figure Table S5**), which together suggested a failure to generate robust polyfunctional T_H_1 cells in SARS-CoV-2 infection. A similar pattern was also observed in SARS-CoV-2-reactive CD4^+^ T cells from healthy non-exposed subjects (**Figures 2B** and **S2C**) but not for HPIV-or HMPV-reactive CD4^+^ T cells, suggesting the defect in generating polyfunctional T_H_1 cells may be a common feature for coronaviruses, although further studies specifically analyzing HCoV-reactive CD4^+^ T cells in healthy individuals will be required to verify this.

Other clusters that were relatively depleted of SARS-CoV-2-reactive CD4^+^ T cells included clusters 9 and 2, which were both enriched for T_H_17 signature genes, with cluster 9 highly enriched for cells expressing *IL17A* and *IL17F* transcripts, thus representing *bona fide* T_H_17 cells (**Figures 2B-2F, S2C-S2E** and **Table S4**). T_H_17 cells have been associated with protective immune responses in certain models of viral infections (Acharya et al., 2017; Wang et al., 2011), however, in other contexts they have been shown to promote viral disease pathogenesis (Ma et al., 2019). Therefore, the functional relevance of an impaired T_H_17 response in COVID-19 is not clear and requires further investigation.

Clusters that were evenly distributed across all viral-specific CD4^+^ T cells include cluster 5 and 3. Cluster 5 displayed a transcriptional profile consistent with enrichment of interferon-response genes (*IFIT3, IFI44L, ISG15, MX2, OAS1*), and cluster 3 was enriched for *CCR7, IL7R* and *TCF7* transcripts, likely representing central memory CD4^+^ T cell subset (**Figures 2B-F, S2C-S2E** and **Table S4**).

Clusters 0,6 and 7, which were colocalized in the uniform manifold approximation and projection (UMAP) plot, were dominated by SARS-CoV-2-reactive CD4^+^ T cells (**Figure 2B**). Cells in these clusters were uniformly enriched for transcripts encoding for cytokines, surface markers and transcriptional coactivators associated with T follicular helper (T_FH_) cell function (*CXCL13, IL21, CD200, BTLA* and *POU2AF1*) (Locci et al., 2013) (**Figures 2B-2F, S2C-S2E** and **Table S4)**. Independent gene set enrichment analysis (GSEA) showed significant positive enrichment of T_FH_ signature genes in these clusters, confirming that cells in these clusters represent circulating T_FH_ cells (**Figure S2G**). *Bona fide* T_FH_ cells reside in the germinal center, however, T_FH_ cells have been described in the blood where increased numbers have been reported in viral infections and following vaccinations (Bentebibel et al., 2013; Koutsakos et al., 2018; Smits et al., 2020). Thus, the increase in circulating SARS-CoV-2-reactive T_FH_ subsets observed in patients with COVID-19 is consistent with published reports in acute infections.

Cluster 12, which expressed high levels of transcripts linked to cell cycle genes *MKI67* and *CDK1*, also contained a large proportion of SARS-CoV-2 reactive CD4^+^ T cells (**Figures 2B-D**), indicative of actively proliferating cells responsive to SARS-CoV-2 antigens. Cluster 4, also dominated by SARS-CoV-2-reactive CD4^+^ T cells, was characterized by high levels of *PRF1, GZMB, GZMH, GNLY* and *NKG7* transcripts, which encode for molecules linked to cytotoxicity (Patil et al., 2018) (**Figures 2B-2F, S2C-S2E** and **Table S4**). GSEA analysis showed significant positive enrichment of signature genes for cytotoxicity in clusters 4 and 8 (**Figure S2G**), confirming these clusters represent cytotoxic CD4^+^ T cells (CD4-CTLs). Overall, our single-cell transcriptomic analysis revealed substantial differences in the nature of CD4^+^ T cell responses to viral infections and highlight subsets that are specifically enriched or depleted in COVID-19 illness.

### SARS-CoV-2-reactive CD4^+^ T cell subsets associated with disease severity

We next assessed if the proportions of SARS-CoV-2 reactive CD4^+^ T cells in any cluster were greater or lower in patients with severe COVID-19 (n=22, requiring hospitalization) when compared to those with milder disease (n=10, not needing hospitalization). Among the three T_FH_ clusters (clusters 0,6 and 7), which consisted almost exclusively of CD4^+^ T cells reactive to SARS-CoV-2, the relative proportion of cells in T_FH_ cluster 6 was greater in patients with severe disease compared to mild disease (**Figures 3A** and **3SA**). Transcripts encoding for transcription factors ZBED2 and ZBTB32 were enriched in the T_FH_ cluster 6 and were also expressed at significantly higher levels in patients with severe disease (**Figures 3B, S3B** and **Table S6**). ZBTB32, also known as PLZP, belongs to a BTB-ZF family of transcriptional repressors like PLZF, BCL6 and ThPOK, has been shown to play a role in impairing anti-viral immune responses by negatively regulating T cell proliferation, cytokine production and development of long-term memory cells (Piazza et al., 2004; Shin et al., 2017). ZBED2, a novel zinc finger transcription factor without a mouse orthologue, has been linked to T cell dysfunction in the context of anti-tumor immune response (Li et al., 2019), and more recently shown to repress expression of interferon target genes (Somerville et al., 2020). In support of potential dysfunctional properties of the cells in the T_FH_ cluster 6, we found increased expression of several transcripts encoding for molecules linked to inhibitory function, like TIGIT, LAG3, TIM3 and PD1 (Thommen and Schumacher, 2018), and to negative regulation of T cell activation and proliferation, like DUSP4 and CD70 (Huang et al., 2012; O’Neill et al., 2017) (**Figures 3B** and **S3C**). Moreover, T_FH_ cells in cluster 6 also expressed high levels of cytotoxicity-associated transcripts (*PRF1, GZMB*) (**Figure 3C** and **S3D**), reminiscent of the recently described cytotoxic T_FH_ cells, which were shown to directly kill B cells and associated with the pathogenesis of recurrent tonsillitis in children (Dan et al., 2019). Together, these findings suggest that T_FH_ cells in cluster 6, which are increased in severe COVID-19 illness, displayed cytotoxicity features that may impair humoral (B cell) immune responses.

**Figure 3:**
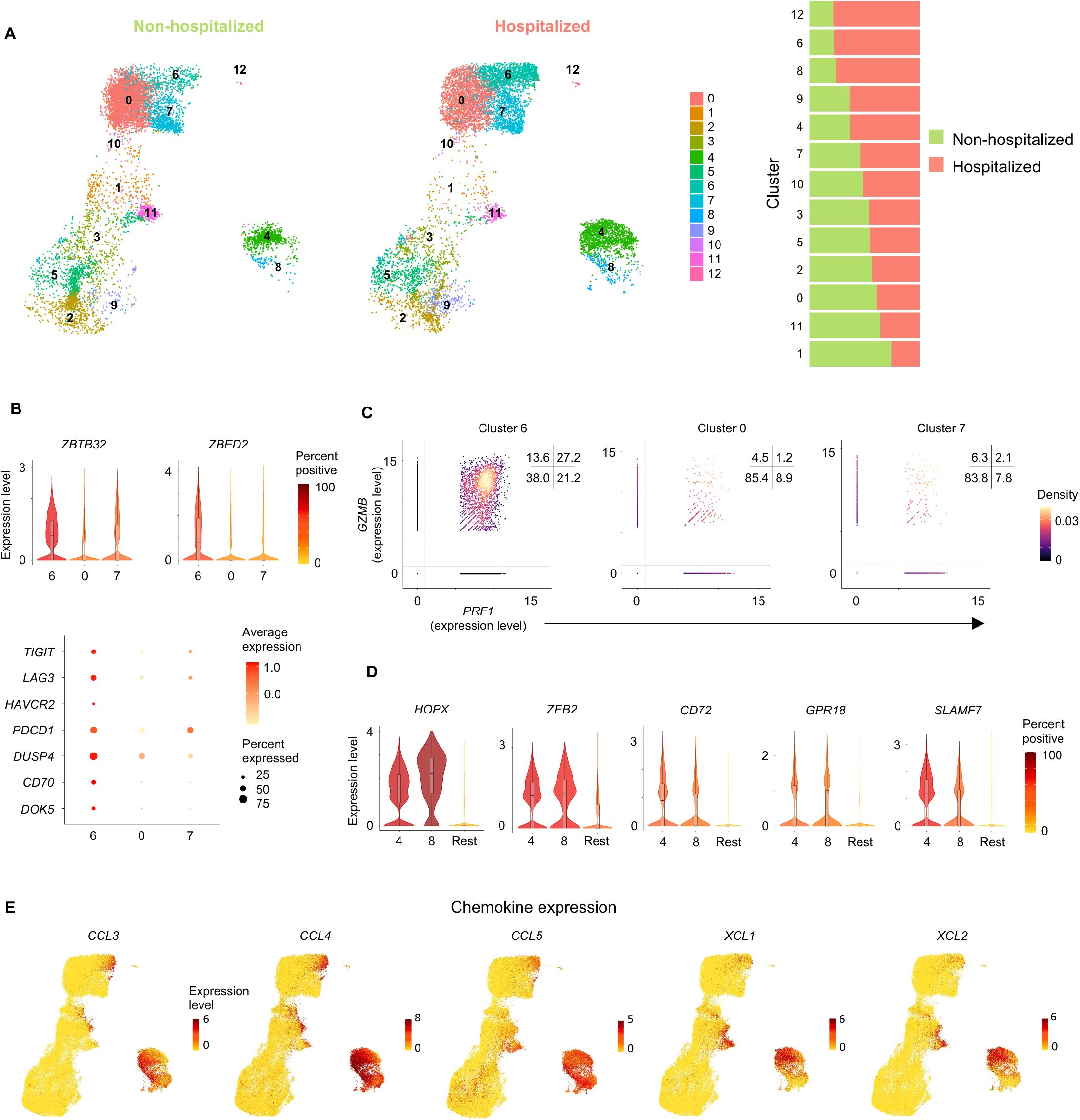
SARS-CoV-2-reactive CD4^+^ T cell subsets associated with disease severity. **(A)** UMAP showing SARS-CoV-2-reactive cells from non-hospitalized and hospitalized COVID-19 patients (left). Normalized proportions of cells per cluster from the two groups (right). **(B)** Violin plots showing expression of *ZBTB32* and *ZBED2* transcripts (left) in SARS-CoV-2-reactive cells from clusters 0,6 and 7 (top); color indicates percentage of cells expressing indicated transcript. Plots below show average expression and percent expression of selected transcripts in indicated clusters. **(C)** Scatter plot displaying co-expression of *PRF1* and *GZMB* transcripts in SARS-CoV-2-reactive cells present in clusters 0,6 and 7. Numbers indicate percentage of cells in each quadrant. **(D)** Violin plots showing expression of *HOPX* and *ZEB2, SLAMF7, CD72* and *GPR18* transcripts in SARS-CoV-2-reactive cells from designated clusters (4, 8) compared to an aggregation of remaining cells (Rest). **(E)** UMAP showing expression of *CCL3, CCL4, CCL5, XCL1* and *XCL2* transcripts in each SARS-CoV-2-reactive cell.

While T cells with cytotoxic function predominantly consist of conventional MHC class I-restricted CD8^+^ T cells, MHC class II-restricted CD4^+^ T cells with cytotoxic potential (CD4-CTLs) have been reported in several viral infections in humans and are associated with better clinical outcomes (Cheroutre and Husain, 2013; Juno et al., 2017; Meckiff et al., 2019; Weiskopf et al., 2015a). Paradoxically, in SARS-CoV-2 infection, we find that cells in the CD4-CTL clusters (cluster 4 and 8) were present at higher frequencies in hospitalized patients with severe disease compared to those with milder disease, potentially contributing to disease severity, although we observed substantial heterogeneity in responses among patients (**Figure 3A** and **Table S3**). Interrogation of the transcripts enriched in the CD4-CTL subsets pointed to several interesting molecules and transcription factors that are likely to play an important role in their maintenance and effector function. These include molecules like CD72 and GPR18 that are known to enhance T cell proliferation and maintenance of mucosal T cell subsets, respectively (Jiang et al., 2017; Wang et al., 2014) (**Figures 3D** and **S3E**). Additional examples include transcription factors HOPX and ZEB2 (**Figures 3D** and **S3E**) that have been shown to positively regulate effector differentiation, function, persistence and survival of T cells (Albrecht et al., 2010; Omilusik et al., 2015). Besides cytotoxicity-associated transcripts, the CD4-CTL subsets (cluster 4 and 8) were highly enriched for transcripts encoding for a number of chemokines like CCL3 (also known as macrophage inflammatory protein (MIP)-1*α*), CCL4 (MIP-1*β*) and CCL5 (**Figures 3E** and **S3F**); these chemokines play an important role in the recruitment of myeloid cells (neutrophils, monocytes, macrophages), NK cells and T cells expressing C-C type chemokine receptors (CCR)1, CCR3 and CCR5 (Hughes and Nibbs, 2018). The CD4-CTL subset in cluster 4 also expressed high levels of transcripts encoding for chemokines XCL1 and XCL2 (**Figures 3E** and **S3G**) that specifically recruit XCR1-expressing conventional type 1 dendritic cells (cDC1) to sites of immune responses where they play a key role in promoting the CD8^+^ T cell responses by antigen cross-presentation (Lei and Takahama, 2012). Overall, the transcriptomic features of SARS-CoV-2-reactive CD4-CTLs suggest that they are likely to be more persistent and play an important role in orchestrating immune responses by recruiting innate immune cells to enhance CD8^+^ T cell responses, while also directly mediating cytotoxic death of MHC class II-expressing virally-infected cells.

### Massive clonal expansion of CD4-CTLs

The recovery of paired T cell receptor (TCR) sequences from individual single cells enabled us to link transcriptome data to clonotype information and evaluate the clonal relationship between different CD4^+^ T cell subsets as well as determine the nature of subsets that display greatest clonal expansion (**Tables S7** and **S8**). In SARS-CoV-2 infection, hospitalized patients were characterized by large clonal expansion of the virus-reactive CD4^+^ T cells (>65%); in contrast, in non-hospitalized patients, less than 45% of TCRs recovered were clonally expanded (**Figure S4A**). Among SARS-CoV-2-reactive CD4^+^ T cells, CD4-CTL subsets (cluster 4 and 8) displayed the greatest clonal expansion (> 75% of cells were clonally-expanded), indicating preferential expansion and persistence of CD4-CTLs in COVID-19 illness (**Figures 4A** and **Table S9**). Analysis of clonally-expanded SARS-CoV-2-reactive CD4^+^ T cells from COVID-19 patients showed extensive sharing of TCRs between cells in clusters 4 and 8, as well as those in cluster 11 (**Figure 4B**), which, notably, was enriched for the expression of *XCL1* and *XCL2* transcripts and also for cytotoxicity-associated transcripts, albeit at lower levels compared to the established CD4-CTL clusters (**Figures 3E, S3G** and **Table S4**). Thus, cells in cluster 11 are likely to be an intermediate transition population, a hypothesis supported by single-cell trajectory analysis that showed potential temporal connection and transcriptional similarity between these subsets (**Figure 4C**).

**Figure 4:**
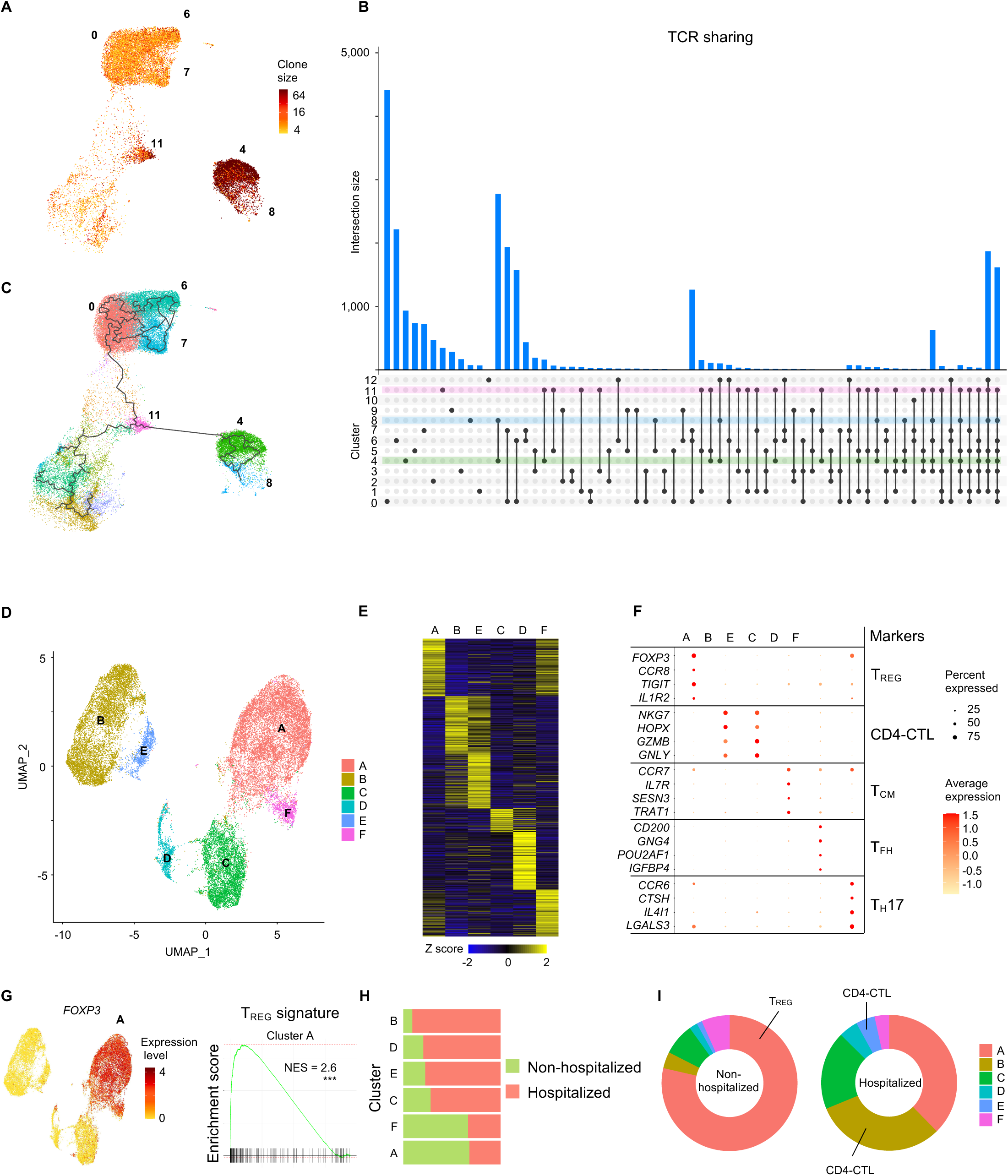
Single-cell TCR sequence analysis, and analysis of SARS-CoV-2-reactive CD4^+^ T cells from 24-hour stimulation condition. **(A)** UMAP showing clone size of SARS-CoV-2-reactive cells from COVID-19 patients (6-hour stimulation condition). **(B)** Single-cell TCR sequence analysis of SARS-CoV-2-reactive cells showing sharing of TCRs between cells form individual clusters (rows, connected by lines). Bars (top) indicate number of cells intersecting indicated clusters (columns). Clusters 4, 8 and 11 are highlighted. **(C)** Single-cell trajectory analysis showing relationship between cells in different clusters (line), constructed using Monocle 3. **(D)** Single-cell transcriptomes of sorted CD137^+^ CD69^+^ memory CD4^+^ T cells following 24 hours stimulation with virus-specific peptide megapools is displayed by UMAP. Seurat-based clustering of 31,341 cells colored based on cluster type. **(E)** Heatmap showing expression of the most significantly enriched transcripts in each cluster (see Table S9, Seurat marker gene analysis – comparison of cluster of interest *versus* all other cells, shown are top 200 transcripts with adjusted P value < 0.05, fold change > 0.25 and >10% difference in the percentage of cells expressing differentially expressed transcript between two groups compared). **(F)** Plot showing average expression and percent expression of selected marker transcripts in each cluster. **(G)** UMAP showing expression of *FOXP3* transcripts (left). GSEA plot for T_REG_ signature genes in cluster A compared to the rest of the cells (right); *** *P* < 0.001. **(H)** Normalized proportions of cells per cluster from non-hospitalized and hospitalized COVID-19 patients is shown. **(I)** Chart showing proportion of SARS-CoV-2-reactive CD4^+^ T cells per cluster from non-hospitalized and hospitalized COVID-19 patients. Notable clusters are highlighted

### SARS-CoV-2-reactive T_REG_ cells are reduced in severe COVID-19 illness

In order to capture SARS-CoV-2-reactive CD4^+^ T cells that may not upregulate the activation markers (CD154 and CD69) after 6 hours of *in vitro* stimulation with SARS-CoV-2 peptide pools, we stimulated PMBCs from the same cultures for a total of 24 hours (see **STAR Methods**) and captured cells based on co-expression of activation markers CD137 (4-1BB) and CD69, a strategy that allowed us to additionally capture antigen-specific regulatory T cells (T_REG_) (Bacher et al., 2016) (**Figures 4D-4G** and **S4B**). Our analysis of a total of 31,278 single-cell CD4^+^ T cell transcriptomes revealed 6 distinct clusters (**Figure 4D-4F** and **Table S9**). The T_FH_ subset (cluster E) was detectable at relatively lower frequencies in the 24-hour condition, though they represented the major CD4^+^ T cell subsets in the 6-hour stimulation condition (**Figures 4D** and **2A**). Consistent with delayed kinetics of activation of central memory T cells (T_CM_ cells), we identified a higher proportion of CD4^+^ T cells expressing transcripts linked to central memory cells (*CCR7, IL7R* and *TCF7*) (cluster C) (**Figures 4D** and **2A**). The largest cluster (cluster A) was characterized by high expression of *FOXP3* transcripts, which encodes for the T_REG_ master transcription factor FOXP3 (**Figures 4D-4G** and **Table S10**). Independent GSEA analysis showed significant positive enrichment of T_REG_ signature genes in this cluster, suggesting that cells in this cluster represented SARS-CoV-2-reactive T_REG_ cells (**Figure 4G**, right). Notably, the T_REG_ cluster contained a relatively lesser proportion of cells from hospitalized COVID-19 patients with severe illness compared to non-hospitalized subjects with milder disease (**Figures 4H** and **4I**), suggesting a potential defect in the generation of immunosuppressive SARS-CoV-2-reactive T_REG_ cells in severe illness. Consistent with our data from 6-hour stimulation condition, we found that cells in the CD4-CTL clusters (cluster B and E) were present at higher frequencies in patients with severe disease (**Figures 4H, 4I** and **S4C**). They also showed the greatest clonal expansion compared to other clusters (**Figure S4D**), suggesting potential importance of the CD4-CTL subset in immune responses to SARS-CoV-2 infection.

## DISCUSSION

There is an urgent need to better understand the molecular determinants of protective and pathogenic immune responses in COVID-19. Given the importance of CD4^+^ T cells in anti-viral immunity, studying this adaptive immune cell population is likely to provide insights into the nature of host responses observed in patients with COVID-19. Current studies on antigen-specific CD4^+^ T cells are limited to flow cytometry-based phenotyping of SARS-CoV-2 responding cells using limited set of markers (Braun et al., 2020; Grifoni et al., 2020; Thieme et al., 2020), which thus fail to comprehensively capture the breath of CD4^+^ T cells that respond to SARS-CoV-2. Unbiased approaches employing single-cell RNA-seq assays can provide these insights, however to our knowledge single-cell studies to-date have only examined total CD4^+^ T cells in blood or bronchoalveolar lavage specimens from patients with COVID-19 illness (Nicolas Vabret et al., 2020). Due to the rarity of SARS-CoV-2-specific cells in the total CD4^+^ T cell populations, signals from these cells are likely to be masked by the relative abundance of other non-antigen specific CD4^+^ T cells. Furthermore, despite a profusion of single-cell transcriptomic studies, the analysis of virus-specific or any antigen-specific T cells, as such in humans has lagged behind, partly due to the challenges imposed by methods to isolate antigen-specific T cells in sufficient numbers. Here, we have overcome these issues and performed one of the largest single-cell transcriptomic study of virus-reactive CD4^+^ T cells, focused on SARS-CoV-2-reactive cells from 32 COVID-19 patients with varying disease severity, and compared their molecular profile to CD4^+^ T cells reactive to other common respiratory viruses.

We find remarkable heterogeneity in the nature of CD4^+^ T cell subsets that are reactive to SARS-CoV-2 and other respiratory viruses, and across patients with differing severity of COVID-19. Polyfunctional T_H_1 cells, which are abundant among FLU-reactive CD4^+^ T cells and are considered to be protective (Seder et al., 2008), were present in lower frequencies among SARS-CoV-2-reactive CD4^+^ T cells. Lower frequencies of T_H_17 cells were also observed among SARS-CoV-2-reactive CD4^+^ T cells. In contrast, we find increased proportions of SARS-CoV-2-reactive T_FH_ cells with dysfunctional and cytotoxicity features in hospitalized patients with severe COVID-19 illness. These findings raise the possibility that certain aspects of antigen-specific CD4^+^ T cell responses required for immune-protection are not optimally generated in COVID-19. Another striking observation is the abundance of CD4-CTLs that express high levels of transcripts encoding for multiple chemokines (XCL1, XCL2, CCL3, CCL4, CCL5) in SARS-CoV-2-reactive CD4^+^ T cells, particularly, from patients with severe COVID-19 illness, suggesting that the CD4-CTL responses in COVID-19 illness may be linked to pathogenesis, although further studies in animal models and large-scale association studies in COVID-19 patients are required to verify or refute this hypothesis. Future studies in COVID-19 patients should also examine the relationships between the subsets of SARS-CoV-2-reactive CD4^+^ T cells in the blood and those observed in the airways and lung tissue where control of SARS-CoV-2 pathogenesis is important.

## STAR METHODS

### CONTACT FOR REAGENT AND RESOURCE SHARING

Further information and requests for reagents may be directed to the corresponding author/lead contacts, Pandurangan Vijayanand and C.H.O (cho.soton.ac.uk).

### EXPERIMENTAL MODEL AND SUBJECT DETAILS

#### COVID-19 patients and samples

Ethical approval for this study from the Berkshire Research Ethics Committee 20/SC/0155 and the Ethics Committee of La Jolla Institute for Immunology (LJI) was in place. Written consent was obtained from all subjects. 22 hospitalized patients in a large teaching hospital in the south of England with SARS-CoV-2 infection, confirmed by reverse transcriptase polymerase chain reaction (RT-PCR) assay for detecting SARS-CoV-2, between April-May 2020 were recruited to the study. A further cohort of 10 participants consisting of healthcare workers who were not hospitalized with COVID-19 illness, confirmed based on RT-PCR assay or serological evidence of SARS-CoV-2 antibodies, were also recruited over the same period. All subjects provided up to 80 mls of blood for research studies. Clinical and demographic data were collected from patient records for hospitalized patients including comorbidities, blood results, drug intervention, radiological involvement, thrombotic events, microbiology and virology results (**Table S1**). The median age of patients with COVID-19 illness was 53 (26-82) and 67% were male. This cohort consisted of 24 (81%) White British/White Other, 4 (13%) Indian and 2 (7%) Black British participants. Of the 32 participants, 10 (30%) had mild disease and were not hospitalized, 22 (70%) had moderate/severe disease and were hospitalized. The median age of the non-hospitalized group was 40 (26-50) and 44% were male. The median age of the hospitalized patients was 60 (33-82) and 76% were male. All hospitalized patients survived to discharge from hospital.

#### Healthy controls

To study HPIV, HMPV and SARS-CoV-2-reactive CD4^+^ T cells from healthy non-exposed subjects (pre-COVID-19 pandemic), we utilized de-identified buffy coat samples from healthy adult donors who donated blood at the San Diego Blood Bank before 2019, prior to the Covid-19 pandemic. Donors were considered to be in good health, free of cold or flu-like symptoms and with no history of Hepatitis B or Hepatitis C infection. To study FLU-reactive cells, we obtained de-identified blood samples from 8 donors enrolled in LJI’s Normal Blood Donor Program before and/or after (12 - 14 days) receiving the FLUCELVAX vaccine. Approval for the use of this material was obtained from the LJI Ethics Committee.

## METHOD DETAILS

### PBMC processing

Peripheral blood mononuclear cells (PBMCs) were isolated from up to 80ml of anti-coagulated blood by density centrifugation over Lymphoprep (Axis-Shield PoC AS, Oslo, Norway) and cryopreserved in 50% decomplemented human antibody serum, 40% complete RMPI 1640 medium and 10% DMSO.

### SARS-CoV-2 peptide pools

Pools of lyophilized peptides covering the immunodominant sequence of the spike glycoprotein and the complete sequence of the membrane glycoprotein of SARS-CoV-2 (15-mer sequences with 11 amino acids overlap) were obtained from Miltenyi Biotec (Thieme et al., 2020) resuspended and stored according to the manufacturer’s instructions.

### SARS-CoV-2 antibody testing

The LIAISON SARS-CoV-2 S1/S2 IgG (DiaSorin S.p.A., Saluggia, Italy) was utilized as per the manufacturer’s instructions to obtain quantitative antibody results from plasma samples via an indirect chemiluminescence immunoassay (CLIA) in an United Kingdom Accreditation Service (UKAS) diagnostic laboratory at University Hospital Southampton. Sample results were interpreted as positive (≥ 15AU/ml), Equivocal (≥ 12.0 and < 15.0 AU/ml) and negative (<12AU/ml).

### Epitope MegaPool (MP) design

The Human Parainfluenza (HPIV), Metapneumovirus (HMPV) CD4^+^ T cell megapools (MPs) were produced by sequential lyophilization of viral-specific epitopes as previously described (Carrasco Pro et al., 2015; Weiskopf et al., 2015b). Full lists of the viral protein sequences derived from the uniprot database and used for the Parainfluenza and Metapneumovirus MP design are available in **Table S11**. T cell prediction was performed using TepiTool tool, available in IEDB analysis resources (IEDB-AR), applying the 7-allele prediction method and a median cutoff ≤20 (Dhanda et al., 2019; Paul et al., 2015; Paul et al., 2016). For the HA-influenza MP, we selected 177 experimentally defined epitopes, retrieved by querying the IEDB database (www.IEDB.org) on 07/12/19 with search parameters “positive assay only, No B cell assays, No MHC ligand assay, Host: Homo Sapiens and MHC restriction class II”. The list of epitopes was enriched with predicted peptides derived from the HA sequences of the vaccine strains available in 2017-2018 and 2018-2019 (A/Michigan/45/2015(H1N1), B/Brisbane/60/2008,A/Hong_Kong/4801/2014_H3N2, A/Michigan/45/2015(H1N1), A/Alaska/06/2016(H3N2), B/Iowa/06/2017, B/Phuket/3073/2013). The resulting peptides were then clustered using the IEDB cluster 2.0 tool and the IEDB recommended method (cluster-break method) with a 70% cut off for sequence identity applied (Dhanda et al., 2019; Dhanda et al., 2018). Peptides were synthesized as crude material (A&A, San Diego, CA), resuspended in DMSO, pooled according to each MP composition and finally sequentially lyophilized (Carrasco Pro et al., 2015). For screening healthy non-exposed subjects (samples provided before the current pandemic) who cross-react to SARS-CoV-2, we screened 20 healthy non-exposed subjects using SARS-CoV-2 peptide CD4-R and CD4-S pools, as described (Grifoni et al., 2020).

### Antigen-reactive T cell enrichment (ARTE) assay

Enrichment and FACS sorting of virus-reactive CD154^+^CD4^+^ memory T cells following peptide pool stimulation was adapted from Bacher *et al*. 2016 (Bacher et al., 2016). Briefly, PBMCs from each donor, were thawed, washed, plated in 6-well culture plates at a concentration of 5 × 10^6^ cells/ml in 1 ml of serum-free TexMACS medium (Miltenyi Biotec) and left overnight (5 % CO_2_, 37 °C). Cells were stimulated by the addition of individual virus-specific peptide pools (1 µg/ml) for 6 h in the presence of a blocking CD40 antibody (1 µg/ml; Miltenyi Biotec). For subsequent MACS-based enrichment of CD154^+^, cells were sequentially stained with fluorescence-labeled surface antibodies (antibody list in **Table S2**), Cell-hashtag TotalSeq(tm)-C antibody (0.5 µg/condition), and a biotin-conjugated CD154 antibody (clone 5C8; Miltenyi Biotec) followed by anti-biotin microbeads (Miltenyi Biotec). Labelled cells were added to MS columns (Miltenyi Biotec) and positively selected cells (CD154^+^) were eluted and used for FACS sorting of CD154^+^ memory CD4^+^ T cells. The flow-through from the column was collected and re-plated to harvest cells responding 24 h after peptide stimulation. Analogous to enrichment for CD154^+^, CD137-expressing CD4^+^ memory T cells cells were positively selected by staining with biotin-conjugated CD137 antibody (clone REA765; Miltenyi Biotec) followed by anti-biotin MicroBeads and applied to a new MS column. Following elution, enriched populations were immediately sorted using a FACSAria Fusion Cell Sorter (Becton Dickinson) based on dual expression of CD154 and CD69 for 6-hour stimulation condition, and CD137 and CD69 for 24-hour stimulation condition. The gating strategy used for sorting is shown in **Figures S1A** and **S4B**. All flow cytometry data were analyzed using FlowJo software (version 10).

### Cell isolation and single-cell RNA-seq assay (10x platform)

For combined single-cell RNA-seq and TCR-seq assays (10x Genomics), a maximum of 60,000 virus-reactive memory CD4^+^ T cells from up to 8 donors were pooled by sorting into low retention 1.5ml collection tubes, containing 500 µl of a1:1 solution of PBS:FBS supplemented with RNAse inhibitor (1:100). Following sorting, ice-cold PBS was added to make up to a volume of 1400 µl. Cells were then centrifuged for 5 minutes (600 g at 4 °C) and the supernatant was carefully removed leaving 5 to 10 µl. 25 µl of resuspension buffer (0.22 µm filtered ice-cold PBS supplemented with ultra-pure bovine serum albumin; 0.04 %, Sigma-Aldrich) was added to the tube and the pellet was gently but thoroughly resuspended. Following careful mixing, 33 µl of the cell suspension was transferred to a PCR-tube for processing as per the manufacturer’s instructions (10x Genomics). Briefly, single-cell RNA-sequencing library preparation was performed as per the manufacturer’s recommendations for the 10x Genomics 5’TAG v1.0 chemistry with immune profiling and cell surface protein technology. Both initial amplification of cDNA and library preparation were carried out with 13 cycles of amplification; V(D)J and cell surface protein libraries were generated corresponding to each 5’TAG gene expression library using 9 cycles and 8 cycles of amplification, respectively. Libraries were quantified and pooled according to equivalent molar concentrations and sequenced on Illumina NovaSeq6000 sequencing platform with the following read lengths: read 1 – 101 cycles; read 2 – 101 cycles; and i7 index - 8 cycles.

### Single-cell transcriptome analysis

Reads from single-cell RNA-seq were aligned and collapsed into Unique Molecular Identifiers (UMI) counts using 10x Genomics’ Cell Ranger software (v3.1.0) and mapping to GRCh37 reference (v3.0.0) genome. Hashtag UMI counts for each TotalSeq(tm)-C antibody capture library were generated with the Feature Barcoding Analysis pipeline from Cell Ranger. To demultiplex donors, UMI counts of cell barcodes were first obtained from the raw data output, and only cells with at least 100 UMI for the hastag with the highest UMI counts were considered for donor assignment. Donor identities were inferred by *MULTIseqDemux* (autoThresh = TRUE and maxiter = 10) from Seurat (v3.1.5) using the UMI counts. Each cell barcode was assigned a donor ID, marked as a Doublet, or having a Negative enrichment. Cells with multiple barcodes were re-classified as doublets if the ratio of UMI counts between the top 2 barcodes was less than 3. Cells labeled as Doublet or Negative were removed from downstream analyses. Raw 10x data, from the 6-hour and 24-conditions were independently aggregated using Cell Ranger’s *aggr* function (v3.1.0). The merged data was transferred to the R statistical environment for analysis using the package Seurat (v3.1.5) (Stuart et al., 2019). To further minimize doublets and to eliminate cells with low quality transcriptomes, cells expressing < 800 and > 4400 unique genes, < 1500 and > 20,000 total UMI content, and > 10% of mitochondrial reads were excluded. The summary statistics for all the single-cell transcriptome libraries are provided in **Table S3** and indicate good quality data with no major differences in quality control metrices across multiple batches (**Figure S2A**). This procedure was independently applied for data from CD4^+^ T cells stimulated for 6 hours and 24 hours.

For single-cell transcriptome analysis only genes expressed in at least 0.1% of the cells were included. The transcriptome data was then log-transformed and normalized (by a factor of 10,000) per cell, using default settings in Seurat software. Variable genes with a mean expression greater than 0.01 mean UMI and explaining 25% of the total variance were selected using the Variance Stabilizing Transformation method, as described (Stuart et al., 2019). Transcriptomic data from each cell was then further scaled by regressing the number of UMI-detected and percentage of mitochondrial counts. For data from CD4+ T cells stimulated for 6 hours, principal component analysis was performed using the variable genes, and based on the standard deviation of PCs in the “elbow plot”, the first 38 principal components (PCs) were selected for further analyses. Cells were clustered using the *FindNeighbors* and *FindClusters* functions in Seurat with a resolution of 0.6. The robustness of clustering was independently verified by other clustering methods and by modifying the number of PCs and variable genes utilized for clustering. Analysis of clustering patterns across multiple batches revealed no evidence of strong batch effects (**Figure S2A**). For data from CD4+ T cells stimulated for 24 hours, principal component analysis was performed using the genes explaining 25% of the variance, and the first 16 principal components (PCs) were selected for further analyses. Cells were clustered using the *FindNeighbors* and *FindClusters* functions in Seurat with a resolution of 0.2. Further visualizations of exported normalized data such has “violin” plots were generated using the Seurat package and custom R scripts. Violin shape represents the distribution of cell expressing transcript of interest (based on a Gaussian Kernel density estimation model) and are colored according to the percentage of cells expressing the transcript of interest.

### Single-cell differential gene expression analysis

Pairwise single-cell differential gene expression analysis was performed using the MAST package in R (v1.8.2) (Finak et al., 2015) after conversion of data to log_2_ counts per million (log_2_ CPM + 1). A gene was considered differentially expressed when Benjamini-Hochberg–adjusted *P*-value was < 0.05 and a log2 fold change was more than 0.25. For finding cluster markers (transcripts enriched in a given cluster) the function *FindAllMarkers* from Seurat was used.

### Gene Set Enrichment Analysis and Signature Module Scores

GSEA scores were calculated with the package *fgsea* in R using the signal-to-noise ratio as a metric. Gene sets were limited by minSize = 3 and maxSize = 500. Normalized enrichment scores were presented as * plots. Signature module scores were calculated with *AddModuleScore* function, using default settings in Seurat. Briefly, for each cell, the score is defined by the mean of the signature gene list after the mean expression of an aggregate of control gene lists is subtracted. Control gene lists were sampled (same size as the signature list) from bins created based on the level of expression of the signature gene list. Gene lists used for analysis are provided in **Table S12**.

### Single-cell trajectory analysis

The “branched” trajectory was constructed using Monocle 3 (v0.2.1, default settings) with the number of UMI and percentage of mitochondrial UMI as the model formula, and including the highly variable genes from Seurat for consistency. After setting a single partition for all cells, the cell-trajectory was projected on the PCA and UMAP generated from Seurat analysis. The ‘root’ was selected by the get_earliest_principal_node function provided in the package.

### T cell receptor (TCR) sequence analysis

Reads from single-cell V(D)J TCR sequence enriched libraries were processed with the *vdj* pipeline from Cell Ranger (v3.1.0 and human annotations reference GRCh38, v3.1.0, as recommended). In brief, the V(D)J transcripts were assembled and their annotations were obtained for each independent library. In order to perform combined analysis of single-cell transcriptome and TCR sequence from the same cells, V(D)J libraries were first aggregated using a custom script. Then cell barcode suffixes from these libraries were revised according to the order of their gene expression libraries. Unique clonotypes, as defined by 10x Genomics as a set of productive Complementarity-Determining Region 3 (CDR3) sequences, were identified across all library files and their frequency and proportion (clone statistics) were calculated based on the aggregation result. This procedure was independently applied for data from CD4^+^ T cells stimulated for 6 and 24 hours. Based on the *vdj* aggregation files, barcodes captured by our gene expression data and previously filtered to keep only good quality cells, were annotated with a specific clonotype ID alongside their clone size (number of cells with the same clonotypes in either one or both the TCR alpha and beta chains) statistics (**Tables S7 and S8**). Cells that share clonotype with more than 1 cell were called as clonally expanded (clone size *≥* 2). Clone size for each cell was visualized on UMAP. Sharing of clonotype between cells in different clusters was depicted using the tool UpSetR (Conway et al., 2017).

## QUANTIFICATION AND STATISTICAL ANALYSIS

Processing of data, applied methods and codes are described in the respective section in the STAR Methods. The number of subjects, samples and replicates analyzed, and the statistical test performed are indicated in the figure legends or STAR methods. Statistical analysis for comparison between two groups was assessed with Student’s unpaired two-tailed t-test using GraphPad Prism.

## DATA AND SOFTWARE AVAILABILITY

Scripts are available in our repository on GitHub (https://github.com/vijaybioinfo/COVID19_2020). Sequencing data for this study is currently been deposited into the Gene Expression Omnibus and a reviewer link will be provided as soon as it is available.

## ACKNOWLEDGMENTS

We thank Luke Smith for patient recruitment and sample collection; Callum Dixon, Benjamin Johnson, Lydia Scarlett and Silvia Austin for collection of clinical data; Céline Galloway, Oliver Wood, Katy McCann and Lindsey Chudley for sample processing; Sharon Gilchrist for the illustration. We thank the La Jolla Institute (LJI) Flow Cytometry Core for assisting with cell sorting; the LJI’s Clinical Studies Core for organizing sample collection. We thank Peter Friedmann, Simon Eschweiler, Anusha Preethi Ganesan for providing critical feedback on the manuscript. This work was funded by NIH grants U19AI142742 (P.V., A.S., C.H.O), U19AI118626 (P.V., A.S., G.S.), R01HL114093 (P.V., F.A., G.S.,), R35-GM128938 (F.A), S10RR027366 (BD FACSAria-II), S10OD025052 (Illumina Novaseq6000), the William K. Bowes Jr Foundation (P.V.), and Whittaker foundation (P.V., C.H.O.). Supported by the Wessex Clinical Research Network and National Institute of Health Research UK.

## END NOTES

### AUTHOR CONTRIBUTIONS

B.M., S.J.C., C.H.O., and P.V. conceived the work. B.M., S.J.C, C.H.O., and P.V. designed the study and wrote the manuscript. S.J.C supervised patient identification, recruitment, sample collection and processing. E.P., supervised the analysis of viral PCR and serology tests. A.G. and A.S., provided essential reagents for the isolation of viral-reactive CD4^+^ T cells. B.M., A.K., performed ARTE assay and FACS sorting, and H.S., performed single-cell RNA-sequencing under the supervision of G.S., C.H.O., and P.V. C.R.S., V.F.R, performed bioinformatic analyses under the supervision of G.S., F.A., C.H.O., and P.V.

## SUPPLEMENTARY FIGURE LEGENDS

**Figure S1.**
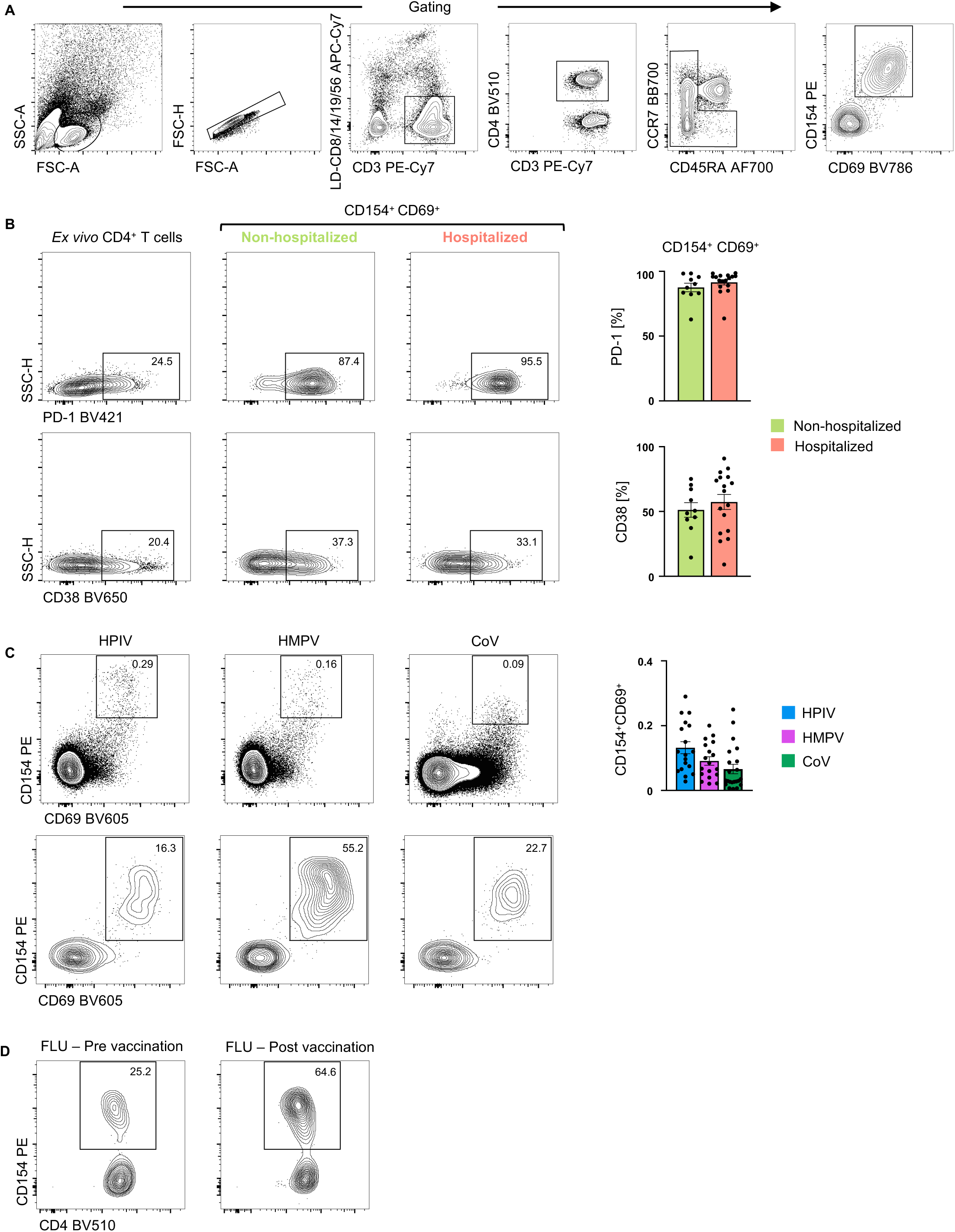
CD4^+^ T cell responses in COVID-19 illness, Related to Figure 1. (A) Gating strategy to sort: lymphocytes size-scatter gate, single cells (Height vs Area forward scatter (FSC)), live, CD3^+^ CD4^+^ memory (CD45RA^+^ CCR7^+^ naïve cells excluded) activated CD154^+^ CD69^+^ cells. Surface expression of activation markers was analyzed on memory CD4^+^ T cells. (B) Representative FACS plots (left) showing surface expression off PD-1 and CD38 in memory CD4^+^ T cells *ex vivo* and in CD154^+^ CD69^+^ memory CD4^+^ T cells following 6 hours of stimulation, post-enrichment (CD154-based); (Right) Percentage of CD154^+^ CD69^+^ memory CD4^+^ T cells expressing PD-1 or CD38 following stimulation and post-enrichment (CD154-based) in 17 hospitalized and 10 non-hospitalized COVID-19 patients; Data are mean +/-S.E.M. (C) Representative FACS plots showing surface staining of CD154 and CD69 in memory CD4^+^ T cells stimulated for 6 hours with individual virus megapools pre-enrichment (top) and post-enrichment (CD154-based) (bottom) in healthy non-exposed subjects. (Right) Percentage of memory CD4^+^ T cells co-expressing CD154^+^ and CD69^+^ following stimulation with individual virus megapools (pre-enrichment); Data are mean +/-S.E.M. (D) Representative FACS plots showing surface staining of CD154 in memory CD4^+^ T cells stimulated with Influenza megapool, post-enrichment (CD154-based), in healthy subjects pre and/or post-vaccination.

**Figure S2.**
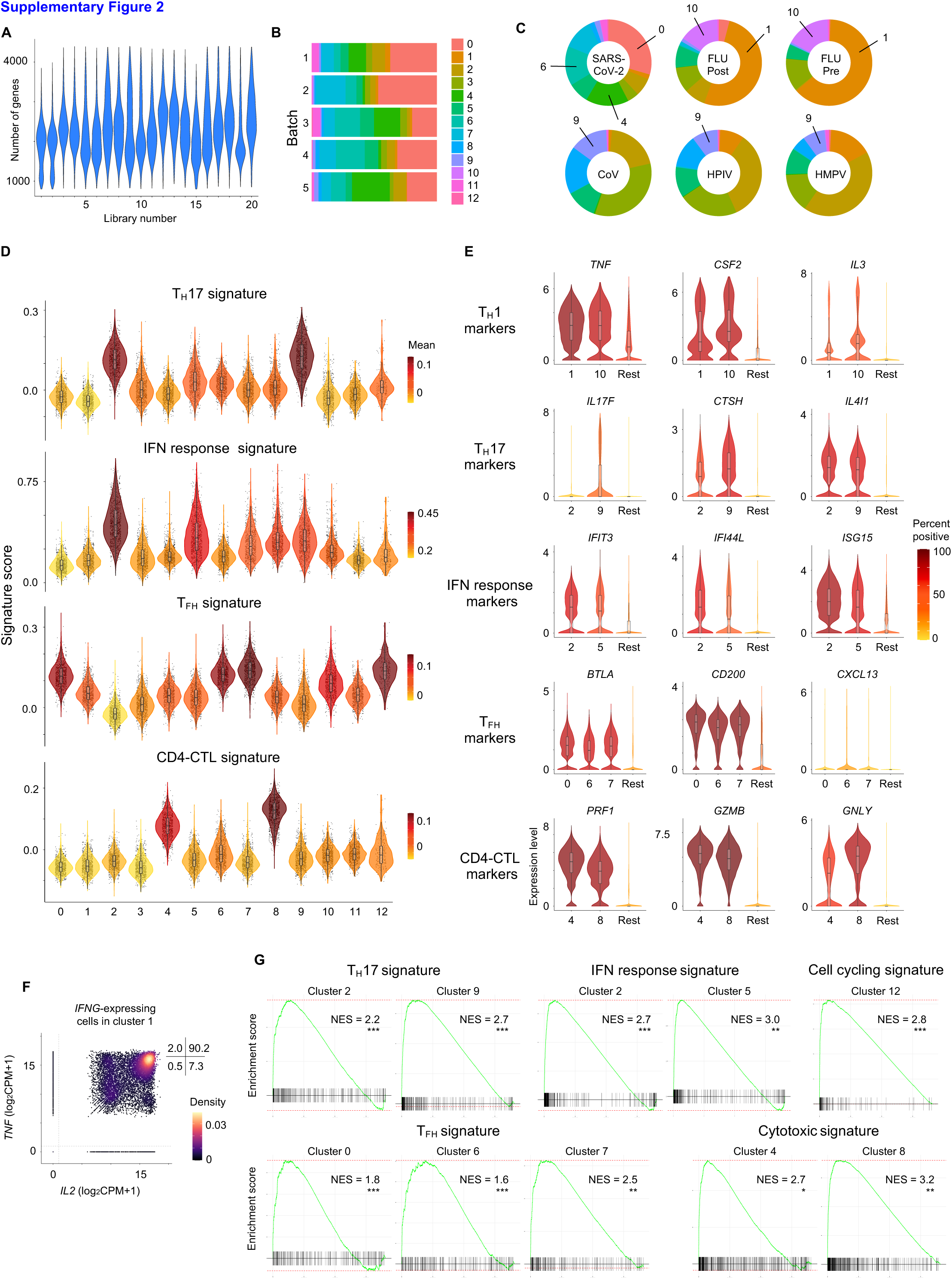
SARS-CoV-2-reactive CD4^+^ T cells are enriched for T_FH_ cells and CD4-CTLs, Related to Figure 2. **(A)** Number of genes recovered for each 10X libraries sequenced. **(B)** Distribution of cells in each cluster for the 5 batches of SARS-CoV-2-reactive CD4^+^ T cells. **(C)** Charts show proportion of individual virus-reactive CD4^+^ T cells per cluster for different viruses. Notable clusters are highlighted. **(D)** Violin plots showing enrichment patterns of T_H_17, interferon (IFN) response, T_FH_, and CD4-CTLs gene signatures for each cluster. Color indicates signature scores. **(E)** Violin plots showing expression of T_H_1, T_H_17, IFN response, T_FH_ and CD4-CTL marker transcripts in designated clusters compared to an aggregation of remaining cells (Rest). Color indicates percentage of cells expressing indicated transcript. **(F)** Scatter plot displaying co-expression of *IL2* and *TNF* transcripts in *IFNG*-expressing, virus-reactive memory CD4^+^ T cells in cluster 1. Numbers indicate percentage of cells in each quadrant. **(G)** Gene set enrichment analysis (GSEA) for T_H_17, cell cycling, T_FH_ and CD4-CTL signature genes in a given cluster compared to the rest of the cells; * *P* < 0.05; *** *P* < 0.01; *** *P* < 0.001.

**Figure S3.**
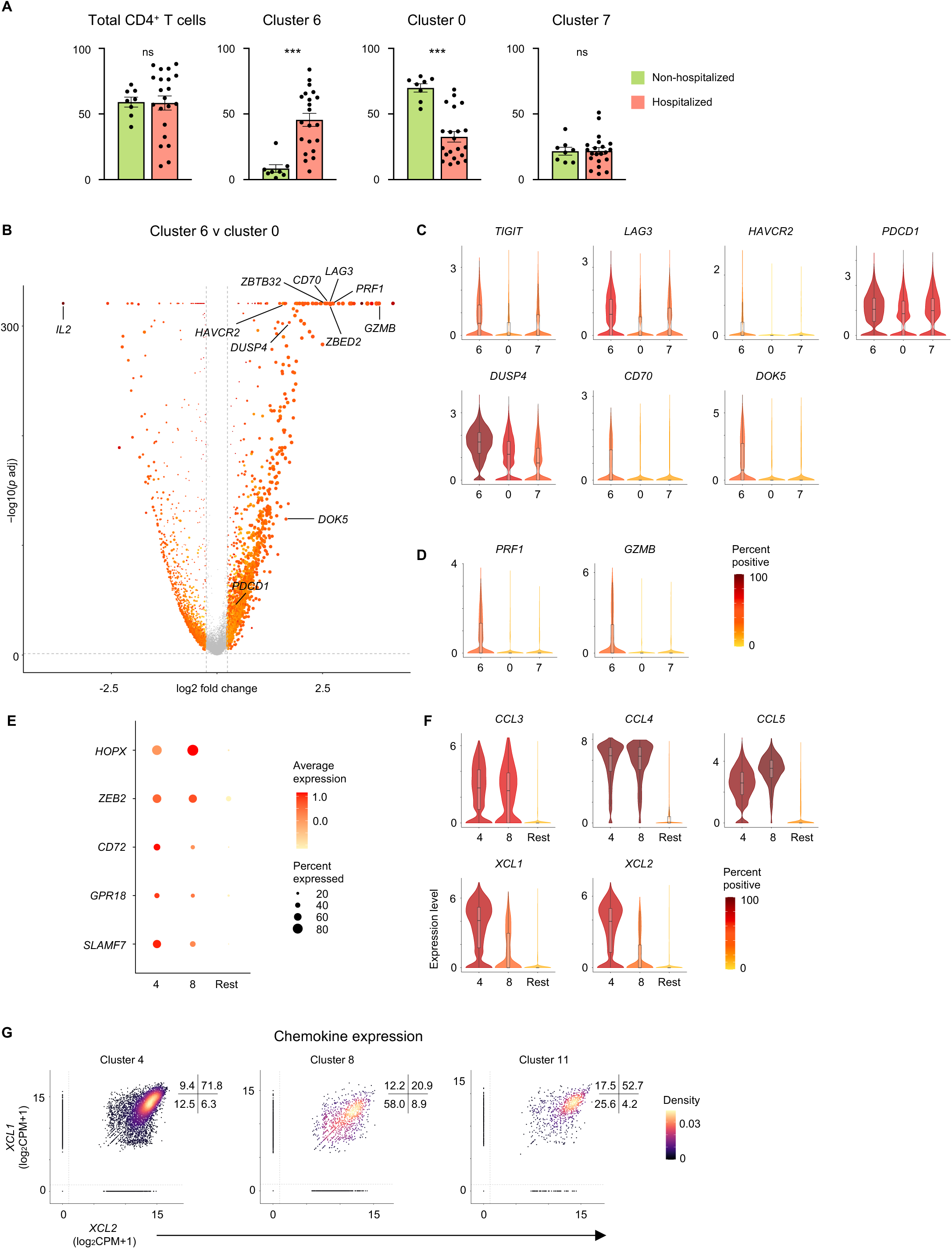
SARS-CoV-2-reactive CD4^+^ T cell subsets associated with disease severity, Related to Figure 3. **(A)** Percentage of T_FH_ cells (clusters 0,6,7) in the total SARS-CoV-2-reactive CD4^+^ T cell pool (left) for non-hospitalized and hospitalized COVID-19 patients; dots indicate data from a single subject (left plot). Other plots show proportion of cluster 6,0,7 cells in SARS-CoV-2-reactive T_FH_ cells in non-hospitalized and hospitalized COVID-19 patient. Data are mean +/-S.E.M; Significance for comparisons were computed using unpaired Student’s t-test (two-tailed); *** *P* < 0.001. **(B)** Volcano plot showing differentially expressed genes between SARS-CoV-2-reactive CD4^+^ T cells in cluster 6 versus cluster 0. **(C)** Violin plots showing expression of *TIGIT, LAG3, HAVCR2, PDCD1, DUSP4, CD70* and *DOK5* transcripts in SARS-CoV-2-reactive cells from clusters 6,0,7 (COVID-19 patients). **(D)** Violin plots showing expression of *PRF1* and *GZMB* transcripts in cells from clusters 6,0,7. **(E)** Average expression and percent expression of selected transcripts in designated clusters (4, 8) compared to an aggregation of remaining cells (Rest). **(F)** Violin plots showing expression *CCL3, CCL4, CCL5, XCL1* and *XCL2* transcripts in designated clusters (4, 8) compared to an aggregation of remaining cells (Rest). **(F)** Scatter plots displaying co-expression of *XCL1* and *XCL2* transcripts in SARS-CoV-2-reactive cells present in designated clusters. Numbers indicate percentage of cells in each quadrant.

**Figure S4.**
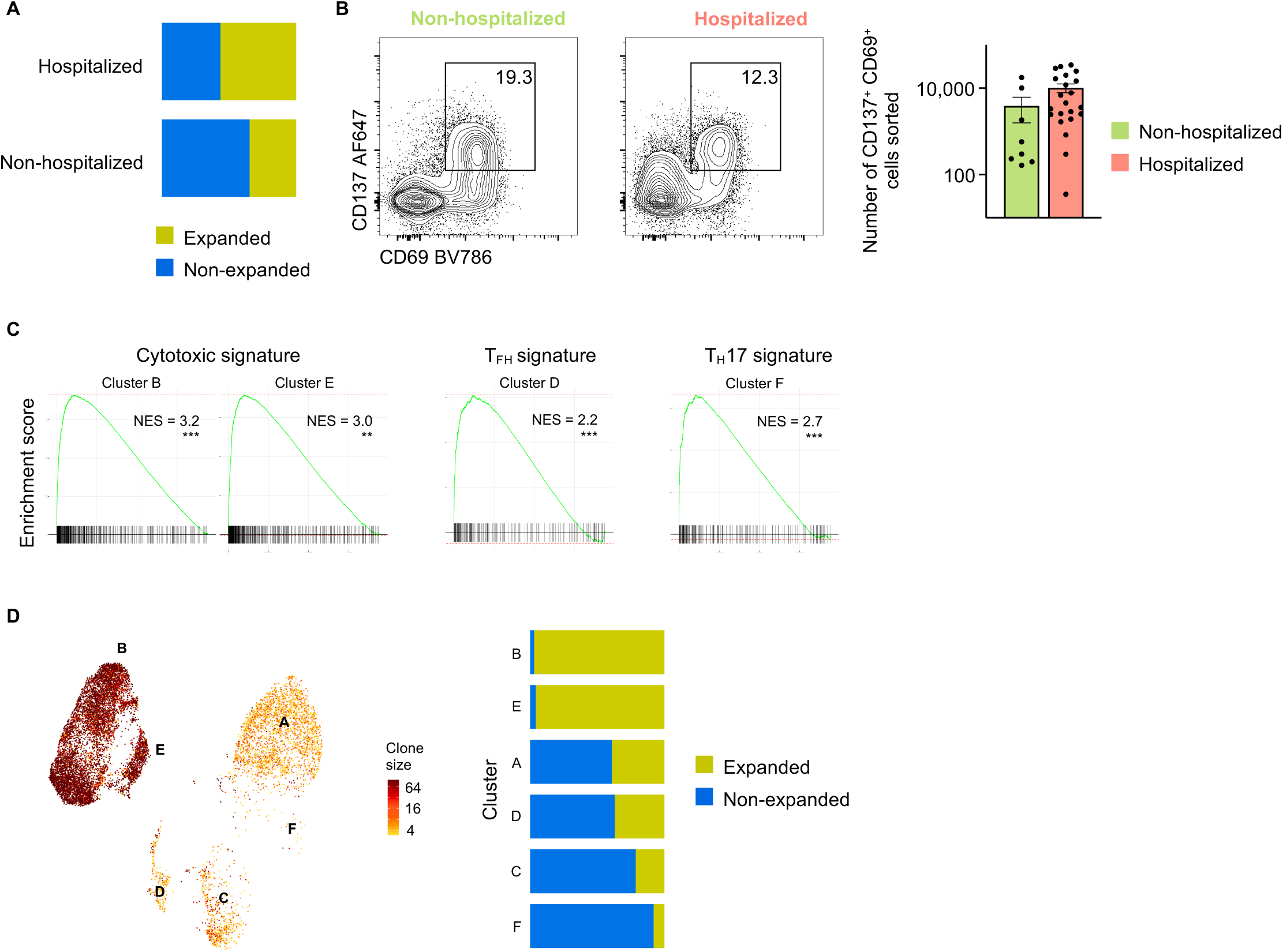
Single-cell TCR sequence analysis, and analysis of SARS-CoV-2-reactive CD4^+^ T cells from 24-hour stimulation condition, Related to Figure 4. **(A)** Proportion of expanded SARS-CoV-2-reactive CD4^+^ T cells (clone size ≥2) in hospitalized and non-hospitalized COVID-19 patients (6-hour stimulation condition). **(B)** Representative FACS plots showing surface staining of CD137 and CD69 in memory CD4^+^ T cells stimulated for 24 hours with SARS-CoV-2 peptide pools, post-enrichment (CD137-based), in hospitalized and non-hospitalized COVID-19 patients (left). Summary of number of cells sorted in 22 hospitalized and 8 non-hospitalized COVID-19 patients (right); Data are mean +/-S.E.M. **(C)** GSEA for cytotoxicity, T_FH_ and T_H_17 signature genes in a given cluster compared to the rest of the cells; *** *P* < 0.001. **(D)** UMAP showing clone size of SARS-CoV-2-reactive cells from COVID-19 patients (24-hour stimulation condition) (left), and proportion of clonally expanded cells (clone size ≥2) in each cluster (right).

## SUPPLEMENTARY TABLE

**Table S1. Human subject details**

**Table S2. Summary of all FACS data**

**Table S3. Single-cell sequencing quality controls and subject-specific cell numbers**

**Table S4. Single-cell cluster enriched transcripts for 6-hour stimulation condition**

**Table S5. Single-cell differential gene expression analysis comparing SARS-CoV-2 versus FLU-reactive CD4**^**+**^ **T cells**

**Table S6. Single-cell differential gene expression analysis**

**Table S7. Subject-specific TCR clonotype data**

**Table S8. Cluster-specific TCR clonotype data**

**Table S9. Single-cell sequencing subject-specific cell numbers**

**Table S10. Single-cell cluster enriched genes for 24-hour stimulation condition**.

**Table S11. Viral protein sequences**

**Table S12. Gene lists utilized for Gene Set Enrichment Analysis and Signature Module Scores**

## REFERENCES

Acharya, D., Wang, P., Paul, A.M., Dai, J., Gate, D., Lowery, J.E., Stokic, D.S., Leis, A.A., Flavell, R.A., Town, T., et al. (2017). Interleukin-17A Promotes CD8+ T Cell Cytotoxicity To Facilitate West Nile Virus Clearance. J Virol 91.

Albrecht, I., Niesner, U., Janke, M., Menning, A., Loddenkemper, C., Kuhl, A.A., Lepenies, I., Lexberg, M.H., Westendorf, K., Hradilkova, K., et al. (2010). Persistence of effector memory Th1 cells is regulated by Hopx. Eur J Immunol 40, 2993–3006.

Bacher, P., Heinrich, F., Stervbo, U., Nienen, M., Vahldieck, M., Iwert, C., Vogt, K., Kollet, J., Babel, N., Sawitzki, B., et al. (2016). Regulatory T Cell Specificity Directs Tolerance versus Allergy against Aeroantigens in Humans. Cell 167, 1067–1078 e1016.

Bacher, P., Hohnstein, T., Beerbaum, E., Rocker, M., Blango, M.G., Kaufmann, S., Rohmel, J., Eschenhagen, P., Grehn, C., Seidel, K., et al. (2019). Human Anti-fungal Th17 Immunity and Pathology Rely on Cross-Reactivity against Candida albicans. Cell 176, 1340–1355 e1315.

Bacher, P., Schink, C., Teutschbein, J., Kniemeyer, O., Assenmacher, M., Brakhage, A.A., and Scheffold, A. (2013). Antigen-reactive T cell enrichment for direct, high-resolution analysis of the human naive and memory Th cell repertoire. J Immunol 190, 3967–3976.

Bentebibel, S.E., Lopez, S., Obermoser, G., Schmitt, N., Mueller, C., Harrod, C., Flano, E., Mejias, A., Albrecht, R.A., Blankenship, D., et al. (2013). Induction of ICOS+CXCR3+CXCR5+ TH cells correlates with antibody responses to influenza vaccination. Science translational medicine 5, 176ra132.

Braun, J., Loyal, L., Frentsch, M., Wendisch, D., Georg, P., Kurth, F., Hippenstiel, S., Dingeldey, M., Kruse, B., Fauchere, F., et al. (2020). Presence of SARS-CoV-2 reactive T cells in COVID-19 patients and healthy donors. medRxiv, 2020.2004.2017.20061440.

Carrasco Pro, S., Sidney, J., Paul, S., Lindestam Arlehamn, C., Weiskopf, D., Peters, B., and Sette, A. (2015). Automatic Generation of Validated Specific Epitope Sets. Journal of immunology research 2015, 763461.

Cheroutre, H., and Husain, M.M. (2013). CD4 CTL: living up to the challenge. Semin Immunol 25, 273–281.

Conway, J.R., Lex, A., and Gehlenborg, N. (2017). UpSetR: an R package for the visualization of intersecting sets and their properties. Bioinformatics 33, 2938–2940.

Dan, J.M., Havenar-Daughton, C., Kendric, K., Al-Kolla, R., Kaushik, K., Rosales, S.L., Anderson, E.L., LaRock, C.N., Vijayanand, P., Seumois, G., et al. (2019). Recurrent group A Streptococcus tonsillitis is an immunosusceptibility disease involving antibody deficiency and aberrant TFH cells. Science translational medicine 11.

Dhanda, S.K., Mahajan, S., Paul, S., Yan, Z., Kim, H., Jespersen, M.C., Jurtz, V., Andreatta, M., Greenbaum, J.A., Marcatili, P., et al. (2019). IEDB-AR: immune epitope database-analysis resource in 2019. Nucleic Acids Res 47, W502–W506.

Dhanda, S.K., Vaughan, K., Schulten, V., Grifoni, A., Weiskopf, D., Sidney, J., Peters, B., and Sette, A. (2018). Development of a novel clustering tool for linear peptide sequences. Immunology.

Finak, G., McDavid, A., Yajima, M., Deng, J., Gersuk, V., Shalek, A.K., Slichter, C.K., Miller, H.W., McElrath, M.J., Prlic, M., et al. (2015). MAST: a flexible statistical framework for assessing transcriptional changes and characterizing heterogeneity in single-cell RNA sequencing data. Genome Biol 16, 278.

Grifoni, A., Weiskopf, D., Ramirez, S.I., Mateus, J., Dan, J.M., Moderbacher, C.R., Rawlings, S.A., Sutherland, A., Premkumar, L., Jadi, R.S., et al. (2020). Targets of T Cell Responses to SARS-CoV-2 Coronavirus in Humans with COVID-19 Disease and Unexposed Individuals. Cell.

Huang, C.Y., Lin, Y.C., Hsiao, W.Y., Liao, F.H., Huang, P.Y., and Tan, T.H. (2012). DUSP4 deficiency enhances CD25 expression and CD4+ T-cell proliferation without impeding T-cell development. Eur J Immunol 42, 476–488.

Hughes, C.E., and Nibbs, R.J.B. (2018). A guide to chemokines and their receptors. FEBS J 285, 2944–2971.

Jiang, X., Bjorkstrom, N.K., and Melum, E. (2017). Intact CD100-CD72 Interaction Necessary for TCR-Induced T Cell Proliferation. Frontiers in immunology 8, 765.

Juno, J.A., van Bockel, D., Kent, S.J., Kelleher, A.D., Zaunders, J.J., and Munier, C.M. (2017). Cytotoxic CD4 T Cells-Friend or Foe during Viral Infection? Frontiers in immunology 8, 19.

Koutsakos, M., Wheatley, A.K., Loh, L., Clemens, E.B., Sant, S., Nussing, S., Fox, A., Chung, A.W., Laurie, K.L., Hurt, A.C., et al. (2018). Circulating TFH cells, serological memory, and tissue compartmentalization shape human influenza-specific B cell immunity. Science translational medicine 10.

Lei, Y., and Takahama, Y. (2012). XCL1 and XCR1 in the immune system. Microbes and infection / Institut Pasteur 14, 262–267.

Li, H., van der Leun, A.M., Yofe, I., Lubling, Y., Gelbard-Solodkin, D., van Akkooi, A.C.J., van den Braber, M., Rozeman, E.A., Haanen, J., Blank, C.U., et al. (2019). Dysfunctional CD8 T Cells Form a Proliferative, Dynamically Regulated Compartment within Human Melanoma. Cell 176, 775–789 e718.

Locci, M., Havenar-Daughton, C., Landais, E., Wu, J., Kroenke, M.A., Arlehamn, C.L., Su, L.F., Cubas, R., Davis, M.M., Sette, A., et al. (2013). Human circulating PD-1+CXCR3-CXCR5+ memory Tfh cells are highly functional and correlate with broadly neutralizing HIV antibody responses. Immunity 39, 758–769.

Ma, W.T., Yao, X.T., Peng, Q., and Chen, D.K. (2019). The protective and pathogenic roles of IL-17 in viral infections: friend or foe? Open Biol 9, 190109.

Meckiff, B.J., Ladell, K., McLaren, J.E., Ryan, G.B., Leese, A.M., James, E.A., Price, D.A., and Long, H.M. (2019). Primary EBV Infection Induces an Acute Wave of Activated Antigen-Specific Cytotoxic CD4(+) T Cells. J Immunol 203, 1276–1287.

Nicolas Vabret, G.J.B., Conor Gruber, Samarth Hegde, Joel Kim, Maria, Kuksin, R.L., Louise Malle, Alvaro Moreira, Matthew D. Park,, Luisanna Pia, E.R., Miriam Saffern, Bérengère Salomé, Myvizhi Esai, Selvan, M.P.S., Jessica Tan, Verena van der Heide, Jill K. Gregory,, Konstantina Alexandropoulos, N.B., Brian D. Brown, Benjamin Greenbaum,, Zeynep H. Gümüş, D.H., Amir Horowitz, Alice O. Kamphorst, Maria A., Curotto de Lafaille, S.M., Miriam Merad, Robert M. Samstein, The, and Project, S.I.R. (2020). Immunology of COVID-19: current state of the science. Immunity (pre-proof) S1074-7613(20)30183-7.

O’Neill, R.E., Du, W., Mohammadpour, H., Alqassim, E., Qiu, J., Chen, G., McCarthy, P.L., Lee, K.P., and Cao, X. (2017). T Cell-Derived CD70 Delivers an Immune Checkpoint Function in Inflammatory T Cell Responses. J Immunol 199, 3700–3710.

Omilusik, K.D., Best, J.A., Yu, B., Goossens, S., Weidemann, A., Nguyen, J.V., Seuntjens, E., Stryjewska, A., Zweier, C., Roychoudhuri, R., et al. (2015). Transcriptional repressor ZEB2 promotes terminal differentiation of CD8+ effector and memory T cell populations during infection. J Exp Med 212, 2027–2039.

Patil, V.S., Madrigal, A., Schmiedel, B.J., Clarke, J., O’Rourke, P., de Silva, A.D., Harris, E., Peters, B., Seumois, G., Weiskopf, D., et al. (2018). Precursors of human CD4(+) cytotoxic T lymphocytes identified by single-cell transcriptome analysis. Science immunology 3.

Paul, S., Lindestam Arlehamn, C.S., Scriba, T.J., Dillon, M.B., Oseroff, C., Hinz, D., McKinney, D.M., Carrasco Pro, S., Sidney, J., Peters, B., et al. (2015). Development and validation of a broad scheme for prediction of HLA class II restricted T cell epitopes. J Immunol Methods 422, 28–34.

Paul, S., Sidney, J., Sette, A., and Peters, B. (2016). TepiTool: A Pipeline for Computational Prediction of T Cell Epitope Candidates. Curr Protoc Immunol 114, 18 19 11–18 19 24.

Piazza, F., Costoya, J.A., Merghoub, T., Hobbs, R.M., and Pandolfi, P.P. (2004). Disruption of PLZP in mice leads to increased T-lymphocyte proliferation, cytokine production, and altered hematopoietic stem cell homeostasis. Mol Cell Biol 24, 10456–10469.

Sallusto, F. (2016). Heterogeneity of Human CD4(+) T Cells Against Microbes. Annu Rev Immunol 34, 317–334.

Seder, R.A., Darrah, P.A., and Roederer, M. (2008). T-cell quality in memory and protection: implications for vaccine design. Nat Rev Immunol 8, 247–258.

Shin, H.M., Kapoor, V.N., Kim, G., Li, P., Kim, H.R., Suresh, M., Kaech, S.M., Wherry, E.J., Selin, L.K., Leonard, W.J., et al. (2017). Transient expression of ZBTB32 in anti-viral CD8+ T cells limits the magnitude of the effector response and the generation of memory. PLoS pathogens 13, e1006544.

Smits, M., Zoldan, K., Ishaque, N., Gu, Z., Jechow, K., Wieland, D., Conrad, C., Eils, R., Fauvelle, C., Baumert, T.F., et al. (2020). Follicular T helper cells shape the HCV-specific CD4+ T cell repertoire after virus elimination. J Clin Invest 130, 998–1009.

Somerville, T.D.D., Xu, Y., Wu, X.S., Maia-Silva, D., Hur, S.K., de Almeida, L.M.N., Preall, J.B., Koo, P.K., and Vakoc, C.R. (2020). ZBED2 is an antagonist of interferon regulatory factor 1 and modifies cell identity in pancreatic cancer. Proc Natl Acad Sci U S A 117, 11471–11482.

Stuart, T., Butler, A., Hoffman, P., Hafemeister, C., Papalexi, E., Mauck, W.M., 3rd, Hao, Y., Stoeckius, M., Smibert, P., and Satija, R. (2019). Comprehensive Integration of Single-Cell Data. Cell 177, 1888–1902 e1821.

Tay, M.Z., Poh, C.M., Renia, L., MacAry, P.A., and Ng, L.F.P. (2020). The trinity of COVID-19: immunity, inflammation and intervention. Nat Rev Immunol.

Thevarajan, I., Nguyen, T.H.O., Koutsakos, M., Druce, J., Caly, L., van de Sandt, C.E., Jia, X., Nicholson, S., Catton, M., Cowie, B., et al. (2020). Breadth of concomitant immune responses prior to patient recovery: a case report of non-severe COVID-19. Nat Med 26, 453–455.

Thieme, C.J., Anft, M., Paniskaki, K., Blazquez-Navarro, A., Doevelaar, A., Seibert, F.S., Hoelzer, B., Konik, M.J., Brenner, T., Tempfer, C., et al. (2020). The SARS-CoV-2 T-cell immunity is directed against the spike, membrane, and nucleocapsid protein and associated with COVID 19 severity. medRxiv, 2020.2005.2013.20100636.

Thommen, D.S., and Schumacher, T.N. (2018). T Cell Dysfunction in Cancer. Cancer Cell 33, 547–562.

Wang, X., Chan, C.C., Yang, M., Deng, J., Poon, V.K., Leung, V.H., Ko, K.H., Zhou, J., Yuen, K.Y., Zheng, B.J., et al. (2011). A critical role of IL-17 in modulating the B-cell response during H5N1 influenza virus infection. Cell Mol Immunol 8, 462–468.

Wang, X., Sumida, H., and Cyster, J.G. (2014). GPR18 is required for a normal CD8alphaalpha intestinal intraepithelial lymphocyte compartment. J Exp Med 211, 2351–2359.

Weiskopf, D., Bangs, D.J., Sidney, J., Kolla, R.V., De Silva, A.D., de Silva, A.M., Crotty, S., Peters, B., and Sette, A. (2015a). Dengue virus infection elicits highly polarized CX3CR1+ cytotoxic CD4+ T cells associated with protective immunity. Proc Natl Acad Sci U S A 112, E4256–4263.

Weiskopf, D., Cerpas, C., Angelo, M.A., Bangs, D.J., Sidney, J., Paul, S., Peters, B., Sanches, F.P., Silvera, C.G., Costa, P.R., et al. (2015b). Human CD8+ T-Cell Responses Against the 4 Dengue Virus Serotypes Are Associated With Distinct Patterns of Protein Targets. J Infect Dis 212, 1743–1751.

